# Beyond resting state neuronal avalanches in the somatosensory barrel cortex

**DOI:** 10.1101/2021.05.13.444047

**Authors:** B. Mariani, G. Nicoletti, M. Bisio, M. Maschietto, R. Oboe, S. Suweis, S. Vassanelli

## Abstract

Since its first experimental signatures, the so called ‘critical brain hypothesis’ has been extensively studied. Yet, its actual foundations remain elusive. According to a widely accepted teleological reasoning, the brain would be poised to a critical state to optimize the mapping of the noisy and ever changing real-world inputs, thus suggesting that primary sensory cortical areas should be critical. We investigated whether a single barrel column of the somatosensory cortex of the anesthetized rat displays a critical behavior. Neuronal avalanches were recorded across all cortical layers in terms of both spikes and population local field potentials, and their behavior during spontaneous activity compared to the one evoked by a controlled single whisker deflection. By applying a maximum likelihood statistical method based on timeseries undersampling to fit the avalanches distributions, we show that neuronal avalanches are power law distributed for both spikes and local field potentials during spontaneous activity, with exponents that are spread along a scaling line. Instead, after the tactile stimulus, activity switches to an across-layers synchronization mode that appears to dominate during cortical representation of the single sensory input.

## 1 INTRODUCTION

The human cortex operates in a state of restless activity, whose meaning and functionality are not yet understood. The critical brain hypothesis suggests that this is the result of the brain operating in the vicinity of the critical point of a phase transition, leading to a rich and variable dynamics at rest. In general, it has been argued that criticality provides biological systems with an optimal balance between robustness against perturbations and the flexibility to adapt to changing conditions. In the case of the brain, this would confer optimal computational capabilities (e.g., by optimizing the correlation length and the dynamic range, leading to the existence of large dynamical repertoires accompanied by maximal transmission and storage of information [1, 2, 3, 4, 5]). In this context, Hidalgo et al. [2] have shown that complex adaptive systems that have to cope with a great variety of stimuli are much more efficient when operating in the vicinity of a critical point, and thus they benefit from dynamically tuning themselves to that point.

By analyzing LFPs of cortical neurons in culture, the seminal work of J. Beggs and D. Plentz [6] provided the first evidence of power law distributed neuronal avalanches, i.e. cascades of activity interspersed by periods of quiescence typical of critical systems. In particular, the exponents of these power laws were remarkably close to the ones of a critical branching process, hence suggesting a scenario in which neuronal networks are characterized by a marginal propagation of the activity at the critical point between an active and an absorbing phase. Since then, such power laws have been observed repeatedly in different experimental settings [7, 8, 9, 10, 11, 12], thus strengthening the critical brain hypothesis. Despite that, ambiguities, inconsistencies and open questions remain.

A first problem concerns the experimental definition of avalanche. In order to estimate avalanches, one needs to define discrete events. While neuronal spikes are events by nature, the conversion of coarse-sampled brain signals such as LFPs into a discrete form (e.g. by simply applying a threshold) is ambiguous and difficult to interpret in terms of neural correlates. As the relation between events and actual underlying neural activity becomes more uncertain, the definition of neural avalanches becomes fuzzy [13, 14, 15]. Moreover, an avalanche should describe a cascade where individual units are *causally* activated, but in experiments causal information is not easily accessible and one needs to resort to various approximation strategies [16]. For example, temporal proximity between events is traditionally considered as a proxy for causality and avalanches are estimated by choosing a discrete time bin Δ*t* that, in turn, depends on the choice of the threshold.

An additional limiting factor is spatial sampling of experimental recordings. Studies using coarse-sampled activity like LFPs typically yielded power-law distributions both in vivo and in vitro, but several experiments relying on spikes in awake animals did not [17, 18, 19]. One possible cause is insufficient spatial sampling of recording in awake conditions. Avalanches are a population phenomenon, but spikes are sparsely recorded in these experiments and reflect a subpopulation of neurons, in fact missing a consistent fraction of the real activity [18]. On the other hand, LFPs are average and composite signals that indeed reflect neuronal populations although they are difficult to interpret in terms of single neurons [20, 21, 22], a problem worsened by the event-extraction process [23].

Also because of these shortcomings, the real nature of the network processes and of the associated phase transition that generate neuronal avalanches has remained elusive. A recent study [24] showed that a statistical meta-analysis of many experiments suggests that the avalanche exponents are not universal, but rather spread along a scaling line. Intriguingly, and in contrast to the classical view of avalanches seen as generated by a quiescent-to-active phase transition, this work suggests that the critical transition in the brain occurs at the edge of synchronization alternatively originating avalanches, oscillations and UP/DOWN states [25].

Up to now, most of the work on neuronal avalanches has focused on spontaneous activity, while much remains to be understood about their behavior after perturbations caused by incoming inputs. Investigating the response to sensory stimuli in primary cortical areas is a clear-cut strategy to address this point. First, these brain regions can be expected to benefit from operating around a critical point to encode the sensory stimuli themselves. Moreover, sensory inputs, which are under direct experimental control, propagate to cortical networks across a limited number of well characterized processing stages (contrary to, e.g., associative or motor areas). Avalanches were studied in the turtle visual cortex ex-vivo while the retina was exposed to a movie acting as a continuous visual stimulus [26, 27]. Based on neural avalanches extracted from LFPs, results suggested that the cortical network self-adapts to a critical state after a short period from the stimulus. However, in these experiments, LFPs were sparsely measured and reflected populations of neurons scattered across the visual processing cascade and downstream to important processing structures including the retina. Work on the primary auditory cortex hinted, instead, at a critical behavior both in resting and post-stimulus conditions [28]. Noteworthy, measurements were confined to either layers 2/3 or 4 and limited by the slow dynamics of calcium imaging to obtain an indirect estimate of neural activity.

In this work we contribute to verify the critical brain hypothesis with a systematic study of neuronal avalanches in the rat barrel cortex (the region of the rat primary somatosensory cortex that encodes tactile sensory inputs from the whiskers). We run our measurements across cortical layers in single barrel columns of the rat anesthetized with tiletamine. This common preparation for electrophysiology [29] displays rich cortical spontaneous activity, including UP and DOWN states and oscillations that have been linked to avalanches and criticality [30, 25]. Given the current challenges to test the critical brain hypothesis and the different results that different kinds of recorded signal can generate [17, 18, 19], we explored activity across a wide frequency range, covering both spikes and LFPs (i.e., up to 3000 Hz). Moreover, analysis was performed both on spontaneous and evoked activity and neural avalanches analyzed through a protocol based on state-of-the-art maximum likelihood statistical method [31].

## 2 MATERIALS AND METHODS

### 2.1 Electrophysiological recordings and surgical procedures

#### Extracellular spikes and LFPs recordings

Spikes were recorded using a neural probe with a linear array of thirty two Iridium Oxide (IrOx) microelectrodes with 65 *µ*m pitch (E32+R-65-S1-L6 NT; Atlas Neuroengineering) (Figure 1). Raw signals were acquired by an Open Ephys Acquisition Board (OEps Tech, Lisbon, Portugal) at 25 KHz sampling frequency and band-pass filtered (300 - 3000 Hz). LFPs were recorded at high density using a CMOS based neural probe with an array of 256 microelectrodes (7.4 *µ*m in diameter size and organized in four vertical columns and sixty-four horizontal rows) [32] with 32 *µ*m pitch along both the horizontal and vertical axis (Figure 1). Raw multiplexed signals were acquired through a NI PXIe-6358 (National Instruments) board (sampling frequency 1.25MS/s at 16bit) and demultiplexed using a home-made LabVIEW software. The resulting whole-array LFP signal was sampled at 976.56 Hz and band-pass filtered (2-300 Hz). Once inserted in the barrel column, both arrays were spanning across all the six cortical layers (from 0 to – 1800 *µ*m).

**Figure 1.**
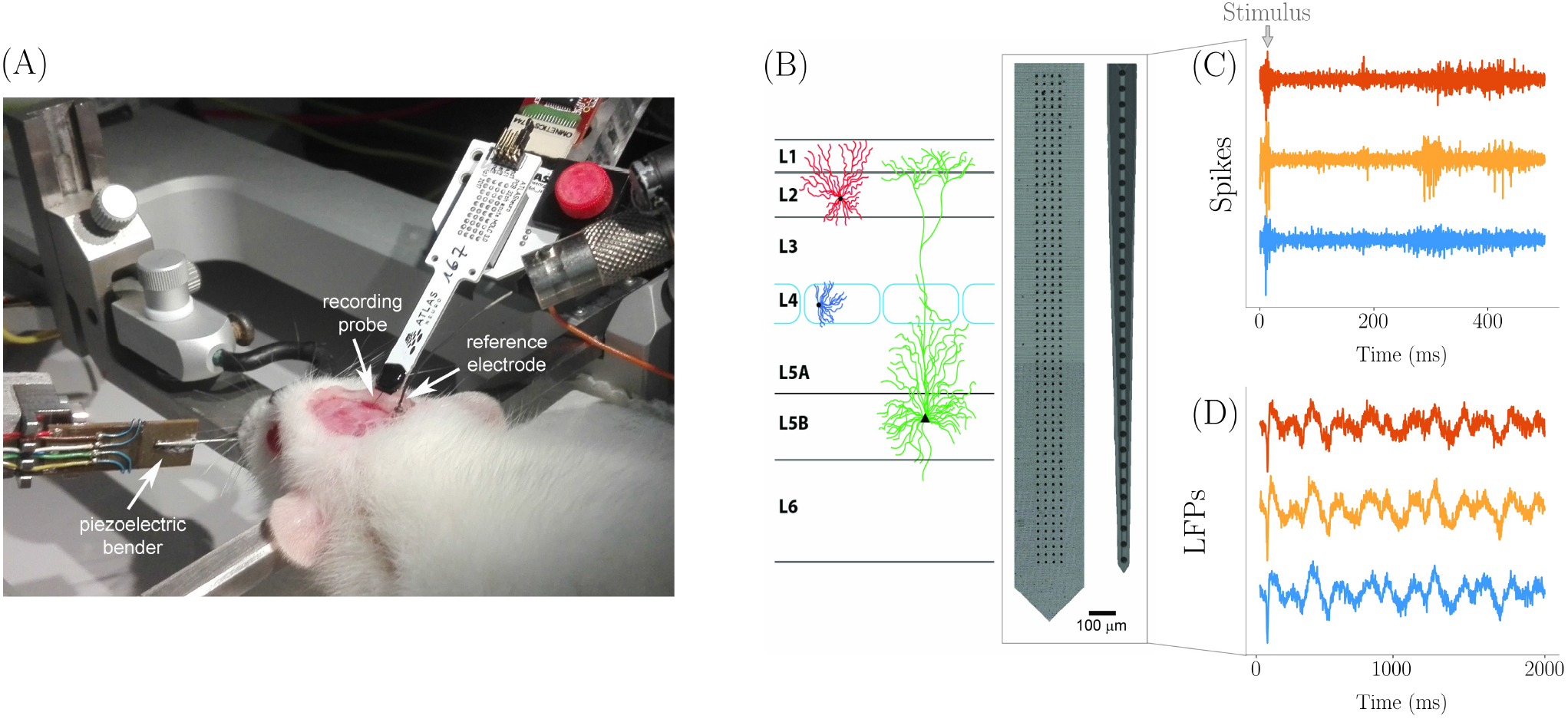
(A) Experimental setting for recording with the anesthetized rat immobilized on the stereotaxic apparatus. The Atlas probe for spikes recording is inserted in the barrel cortex through the dedicated cranial window. The cortical surface is bathed by Krebs’ solution that is grounded through the immersed Ag/AgCl reference electrode. The piezoelectric bender with cannula used to control single whisker deflection is visible on the left. An identical arrangement, but with the custom high-density probe, was adopted for LFPs recording. (B) Custom 2D 64 *×* 4 array used for LFPs and commercial 1D array used for spikes spanning across cortical layers as during recording. The actual number of electrodes inserted in the barrel cortex is 220 for LFPs (organized in a 55 *×* 4 matrix form) and 27 for spikes. Examples of spikes (C) and LFPs (D) traces (from three representative channels each).

#### Surgical implantation and single whisker stimulation

Wistar rats were maintained under standard environmental conditions in the animal research facility of the Department of Biomedical Sciences - University of Padova. All the procedures were approved by the local Animal Care Committee (O.P.B.A.) and the Italian Ministry of Health (authorization number 522/2018-PR). Rats of both genders, aged 36 to 50 days (P36 - P50) and weighting between 150 and 230 g, were anesthetized with an intra-peritoneal induction mixture of tiletamine-xylazine (2 mg and 1.4 g/100 g body weight, respectively), followed by additional doses (0.5 mg and 0.5 g/100 g body weight) every hour. The anesthesia level was constantly monitored by testing the absence of eye and hind-limb reflexes and whiskers’ spontaneous movements. Before starting with surgery, the rat was fixed on a stereotaxic apparatus by teeth and ear bars. The body temperature was monitored continuously with a rectal probe and maintained at 37 °C by a heating pad. The skull was exposed through an anterior-posterior opening of the skin in the center of the head and a window was drilled over the right somatosensory barrel cortex at stereotaxic coordinates – 1 *÷* –4 AP, +4 *÷* +8 ML referred to bregma [33]. A slit in the meninges was made with dedicated fine forceps at coordinates *–*2.5 AP, +6 ML for the subsequent insertion of the recording probe, and the brain was constantly bathed in Krebs’ solution (in mM: NaCl 120, KCl 1.99, NaHCO_3_ 25.56, KH_2_ PO_4_ 136.09, CaCl_2_ 2, MgSO_4_ 1.2, glucose 11).

The recording probe was fixed to a dedicated holder connected to a Patchstar micromanipulator (Scientifica Ltd, East Sussex, UK), which was used for inserting the probe into the cortex orthogonal to the cortical surface. The depth was set at 0 *µ*m when the electrode proximal to the chip tip touched the cortical surface. An Ag/AgCl electrode bathed in Krebs’ solution in proximity of the probe was used as reference.

Contralateral whiskers were trimmed at around 10 mm from the mystacial pad. To control deflection, single whiskers were inserted for 8 mm inside a cannula glued to a piezoelectric bender with integrated strain gauges (P-871.122; Physik Instrumente (PI) GmbH & Co. KG) and driven by a home-made closed-loop control system. Each stimulus, delivered by a waveform generator (Agilent 33250A 80 MHz, Agilent Technologies Inc., Colorado, USA), was consisting of a voltage pulse of 5 ms duration and 100 *µ*s rise/fall time applied to the piezoelectric bender. The principal (maximally responding) whisker identified on the basis of the amplitude of evoked LFP responses was selected for the recording session.

### 2.2 Avalanches analysis

Spikes and high-density LFPs recordings were performed separately in four and five rats, respectively. The minimum time interval between consecutive whisker stimuli was set to two seconds to avoid receptors and central adaptation phenomena. Accordingly, two seconds of recording after the stimuli were excluded from the analysis of *basal* activity. Due to the different duration of the stimulus-evoked avalanches in the LFPs and spikes domains (Fig. 2), *post-stimulus* intervals were set for analysis at 2 s or 500 ms for LFPs and spikes, respectively. In total, 2 minutes long recordings of LFPs basal activity and 5 minutes long recordings of spikes basal activity were analyzed for each rat. Forty stimulations of the whisker were considered for each rat in the analysis of LFPs evoked activity, while in spikes data the recordings of each rat included at least 60 stimulations of the whisker.

**Figure 2.**
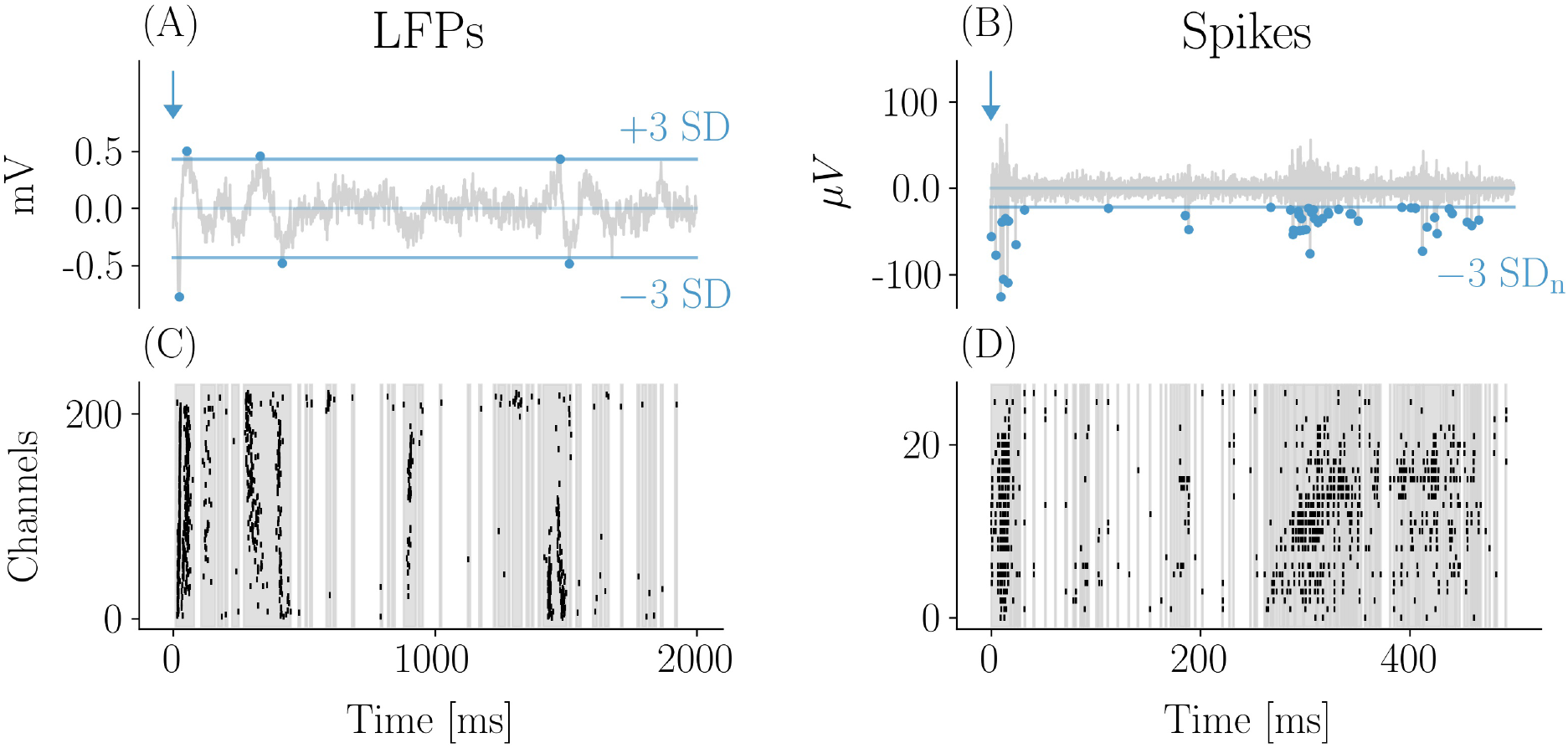
Evoked response by whisker stimulation: LFPs and spikes. Example of LFP (A) and spike (B) traces with the 3 standard deviations (SD) thresholds used for events detection (indicated by dots). The corresponding raster plots are shown in (C) and (D). The 2D array in (C) was rearranged in one dimension (groups of four sensors in a row are reported consecutively along the vertical axis). The early response to the stimulus occurred within few tens of milliseconds and was characterized by a high degree of events synchronization across channels both for LFPs and spikes. The typical LFP evoked response consisted in a first negative peak followed by a slower positive wave. Both were detected as events which is reflected in the raster plot. After the early response large and frequent avalanches typically followed, taking the form of oscillations of synchronous events across channels fading after a few hundreds of milliseconds from the stimulus. Correspondingly, epochs of high frequency firing (bursts) were observed in the spikes domain (C and D).

For the detection of LFP events, the standard deviation (SD) and the mean of the signal was computed for each channel. Both negative and positive deflections of the potential trace were considered *events* when above a threshold of three SD. Moreover, each deflection was considered terminated only after it crossed the mean of the signal.

Noteworthy, in order to distinguish real events from noise, the choice of three SD was based on the distribution of the signal amplitudes which significantly deviated from a Gaussian best fit above that threshold (see Supplementary Material). For both post stimulus and basal (resting state) activity, an average inter event interval (⟨IEI⟩) was calculated and used for temporal binning to estimate avalanches. Avalanches were defined as sequences of ⟨IEI⟩ time bins presenting activity in the form of events, with the end of the avalanche identified by the first empty bin. The number of events in each avalanche accounted for its size, while the duration was the number of temporal bins comprising the avalanche.

For extracellular spikes detection we used as threshold three SD of the noise [34]. Events recorded at the same time frame by different microelectrodes were ascribed to the same neuron and thus counted as one event, although avalanches results were not significantly affected by this correction. Statistics of neuronal avalanches were studied in the four rats, in resting state recordings and in recordings with a stimulation session.

### 2.3 Power law fitting and statistical testing

The avalanches sizes and durations distributions are fitted using the maximum likelihood method. The fitting function for both avalanche sizes and duration is a discrete power-law:

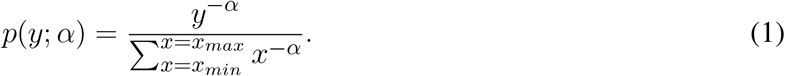

The parameter *x*_*max*_ is set to the maximum observed size or duration. Then the tails of the distributions are fitted by selecting as parameter *x*_*min*_ the one that minimizes the Kolmogorov-Smirnov distance (KS), following the method proposed by Clauset et al. [35]:

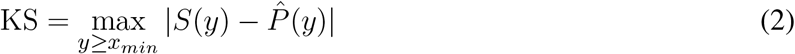

where *S*(*y*) is the cumulative distribution function (CDF) of the data and 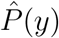 is the CDF of the theoretical distribution fitted with the parameter that best fits the data for *y ≥ x*_*min*_.

After finding the best-fit power law, to assess goodness-of-fit we compared the experimental data against 1000 surrogate datasets drawn from the best-fit power law distribution with the same number of samples as the experimental dataset. The deviation between the surrogate datasets and a perfect power law was quantified with the KS statistic. The p-value of the power-law fit was defined as the fraction of these surrogate KS statistics which were greater than the KS statistic for the experimental data. Note that the data were considered power law distributed if the null hypothesis could not be rejected, namely if the the p-value turned out to be greater than the significance level, which was set to a conservative value of 0.1.

However, when estimating the parameters and evaluating the p-value, we take into consideration another aspect that has been recently pointed out in [31]. A point often ignored is that maximum likelihood methods rely on two assumptions:

1. the observations *y* are distributed as *p*(*y*; *α*), where *α* is the power law exponent;

2. the empirical observations *y*_*i*_, *i* = 1, …, *N*, are independent.

While the first assumption corresponds to our choice of a statistical law, statistical tests rely on the second one, which for instance is implicitly assumed when the log-likelihood is computed as 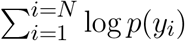. However, complex systems are often characterized by strong temporal and spatial inter-dependencies, thus often violating the independence assumption. This may lead to false rejections of the statistical laws and to over-optimistic uncertainties of the estimated parameters. The authors of [31] propose a method to distinguish between these assumptions, and we exploit it here to estimate the parameters and evaluate the goodness-of-fit. Briefly, we take the timeseries of sizes or durations of consecutive avalanches, and we estimate the time *τ*^*\**^ after which to observations (e.g. the avalanche sizes) are independent from each other. In practice, *τ*^*\**^ is obtained by computing the time at which the autocorrelation of the timeseries reaches an interval around zero (1-percentile of the random realization). Then, the original sequence of length *N* is randomized, *N*^*\**^ = *N/τ*^*\**^ observations are selected and the standard statistical analysis is applied to the new sample. This guarantees that the new sample of dimension *N* ^*\**^ *< N* comprises only uncorrelated avalanches.

Indeed, in our timeseries of sizes and durations we find non negligible values of *τ*^*\**^ (see Supplementary Material), and we verify that the acceptance rate of the statistical law increases for uncorrelated avalanches. Because of the variability of the different realizations of the subsampling procedure, the exponents and the p-values shown in Tables 1 and 2 are obtained averaging over 20 repetitions of this subsampling.

**Table 1.** Exponents before and after subsampling in LFP data, with the corresponding p-values. Note that the p-values obtained after subsampling are always greater than the significance level 0.1 both for *τ* and *τ*_*t*_, except for the *τ*_*t*_ p-value of rat 3.

| | | $\tau$ | p-values | $\tau$ (sub.) | p-values (sub.) | $\tau_t$ | p-values | $\tau_t$ (sub.) | p-values (sub.) |
| --- | --- | --- | --- | --- | --- | --- | --- | --- | --- |
| Rat 1 | Post stim. | 1.85 | 0.04 | 1.87 | 0.48 | 2.44 | 0.16 | 2.45 | 0.50 |
|  | Resting state | 1.82 | < 0.001 | 1.80 | 0.33 | 2.38 | 0.40 | 2.36 | 0.63 |
| Rat 2 | Post stim. | 1.59 | 0.1 | 1.59 | 0.50 | 1.87 | 0.004 | 1.88 | 0.14 |
|  | Resting state | 1.60 | 0.02 | 1.60 | 0.38 | 1.84 | < 0.001 | 1.84 | 0.24 |
| Rat 3 | Post stim. | 1.60 | 0.13 | 1.60 | 0.41 | 2.19 | 0.05 | 2.18 | 0.36 |
|  | Resting state | 1.57 | 0.07 | 1.58 | 0.60 | 1.87 | < 0.001 | 1.87 | 0.03 |
| Rat 4 | Post stim. | 1.98 | 0.27 | 1.97 | 0.56 | 2.78 | 0.41 | 2.75 | 0.7 |
|  | Resting state | 2.16 | 0.54 | 2.15 | 0.72 | 2.67 | 0.30 | 2.65 | 0.58 |
| Rat 5 | Post stim. | 1.74 | 0.30 | 1.73 | 0.71 | 2.19 | 0.07 | 2.19 | 0.45 |
|  | Resting state | 1.72 | 0.56 | 1.73 | 0.78 | 2.18 | 0.09 | 2.20 | 0.55 |

**Table 2.** Exponents before ad after subsampling in spikes data, with the corresponding p-values. Note that the p-values obtained after subsampling are always greater than the significance level 0.1 both for *τ* and *τ*_*t*_, except for the *τ*_*t*_ p-value of rat 4.

| | | $\tau$ | p-values | $\tau$ (sub.) | p-values (sub.) | $\tau_t$ | p-values | $\tau_t$ (sub.) | p-values (sub.) |
| --- | --- | --- | --- | --- | --- | --- | --- | --- | --- |
| Rat 1 | Post stim. | 1.98 | 0.077 | 1.97 | 0.63 | 2.27 | 0.27 | 2.30 | 0.79 |
|  | Resting state | 2.23 | 0.2 | 2.23 | 0.66 | 2.52 | 0.34 | 2.52 | 0.67 |
| Rat 2 | Post stim. | 2.00 | 0.73 | 1.95 | 0.74 | 2.25 | 0.41 | 2.27 | 0.74 |
|  | Resting state | 2.04 | 0.003 | 2.03 | 0.43 | 2.23 | < 0.001 | 2.23 | 0.1 |
| Rat 3 | Post stim. | 2.05 | 0.72 | 2.04 | 0.75 | 2.34 | 0.84 | 2.29 | 0.73 |
|  | Resting state | 2.18 | 0.004 | 2.18 | 0.42 | 2.41 | < 0.001 | 2.41 | 0.26 |
| Rat 4 | Post stim. | 1.98 | 0.06 | 2.00 | 0.91 | 2.51 | 0.01 | 2.33 | 0.63 |
|  | Resting state | 2.50 | 0.12 | 2.50 | 0.19 | 2.67 | < 0.001 | 2.60 | < 0.001 |

## 3 RESULTS

### 3.1 Preliminary considerations

The standard approach to infer criticality is to search for neuronal avalanches whose sizes and durations follow scale-free distributions in resting state (i.e., unperturbed) conditions. Indeed, at criticality, it is expected that such distributions scale as the power laws

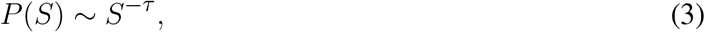

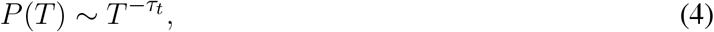

where *S* is the number of events in an avalanche (i.e., the size), *T* is its duration (also called avalanche lifetime) and *τ* and *τ*_*t*_ are the related critical exponents. Power laws, however, can also stem from non critical systems and generative mechanisms. Thus, a more robust test of criticality is to verify whether the so-called crackling noise relation holds. This scaling relation was first developed in the context of crackling noise [36], hence the name, but nonetheless it is expected to hold in general in all systems close to their critical point [37], and in particular in systems with absorbing states [38]. The relation predicts that the critical exponent *δ*, which relates the duration of an avalanche to its mean size as

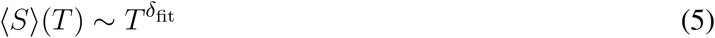

obeys the scaling relation

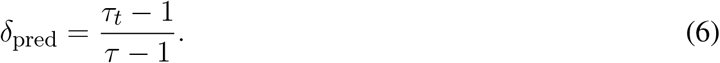

We estimated *δ* in two independent ways, as *δ*_pred_ and as *δ*_fit_, i.e., the slope of the least square fit of the average sizes given their durations. In principle, if these two estimates are compatible, then the system is compatible with criticality. Proving such relation is however challenging. First, it is sensitive to the fitting methods of the distributions of avalanche sizes and lifetimes, as it has been recently shown [39]. Second, in the case of LFPs, the range of avalanche lifetimes typically extends over one order of magnitude only, which undermines reliability of power law fitting.

A complementary approach is to look for criticality by perturbing the system, that is by shifting activity to a subcritical or supercritical regime and then measure the distance from a critical state [5, 40]. Thus, when measuring neural activity across a cortical barrel column of the rat brain, we also provided sensory stimuli consisting of impulsive deflections of the corresponding whisker and therefore representing strong, well defined, and accurately reproducible perturbations.

Neural avalanches sizes and durations were fitted with a discrete power law through the maximum likelihood method (see Section 2.3). As mentioned in the Materials and Methods section, we found non negligible values of the correlation time *τ*^*\**^ both for spikes and LFPs, suggesting that events were not independent. Therefore, to test if avalanches were power law distributed, we corrected for dependencies between avalanches by subsampling the data of sizes and durations as described in Section 2.3.

### 3.2 Avalanches in LFP Data

We first analyzed avalanches in LFP recordings and found similar results across five animals (Table 1). First of all, we focused on spontaneous (i.e., resting, non stimulated) activity. We estimated *τ* ^*\**^ and subsampled the sizes and the lifetimes accordingly to ensure that individual observations were uncorrelated (thus not violating the assumption of the maximum likelihood method). Following this correction, avalanches resulted power law distributed across the five rats, except for one case for the avalanches durations (see Figure 4 and Table 1).

**Figure 3.**
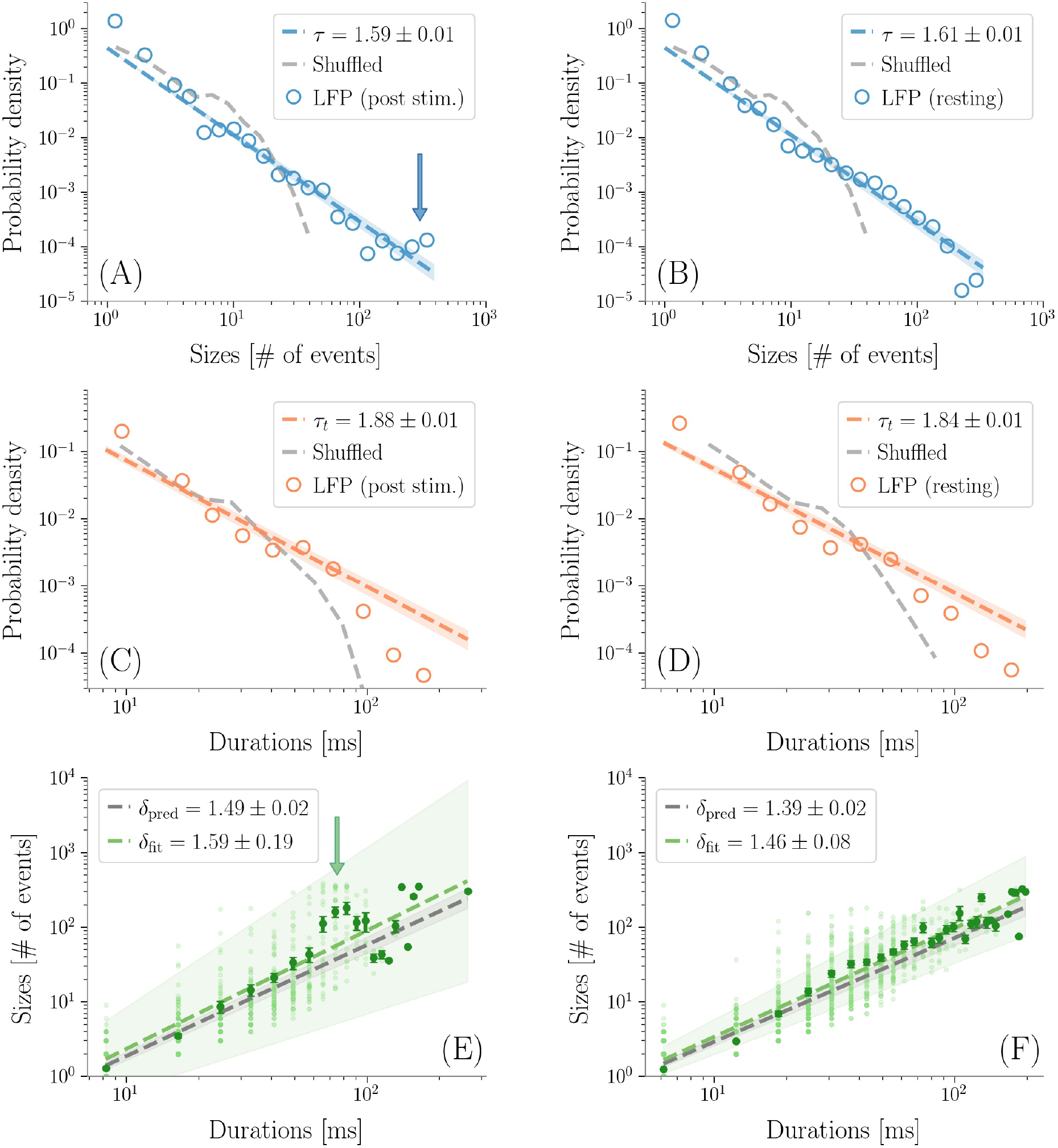
LFP avalanches. Representative probability densities of avalanches sizes in one rat after the sensory stimulus (A), or in resting state conditions (B). A bump deviating from the power law can be recognized in the post-stimulus distribution in correspondence of large avalanche sizes, pointed to by an arrow. Distributions of the avalanches’ durations do not display clear alterations instead (C-D). Durations are expressed in *ms* by multiplying the number of bins by ⟨IEI⟩. In the randomized dataset the exponential distribution provides a better fit in all cases (dashed gray lines). The shuffling procedure consists in randomizing the occurrence times of the events of each channel, so that the events rate of each channel is preserved. (E-F) The crackling noise relation is verified in both cases within the experimental errors (i.e., *δ*_pred_ is compatible with *δ*_fit_). Once more, notice in (E) the presence of a bump (indicated by an arrow) in the post-stimulus regime.

**Figure 4.**
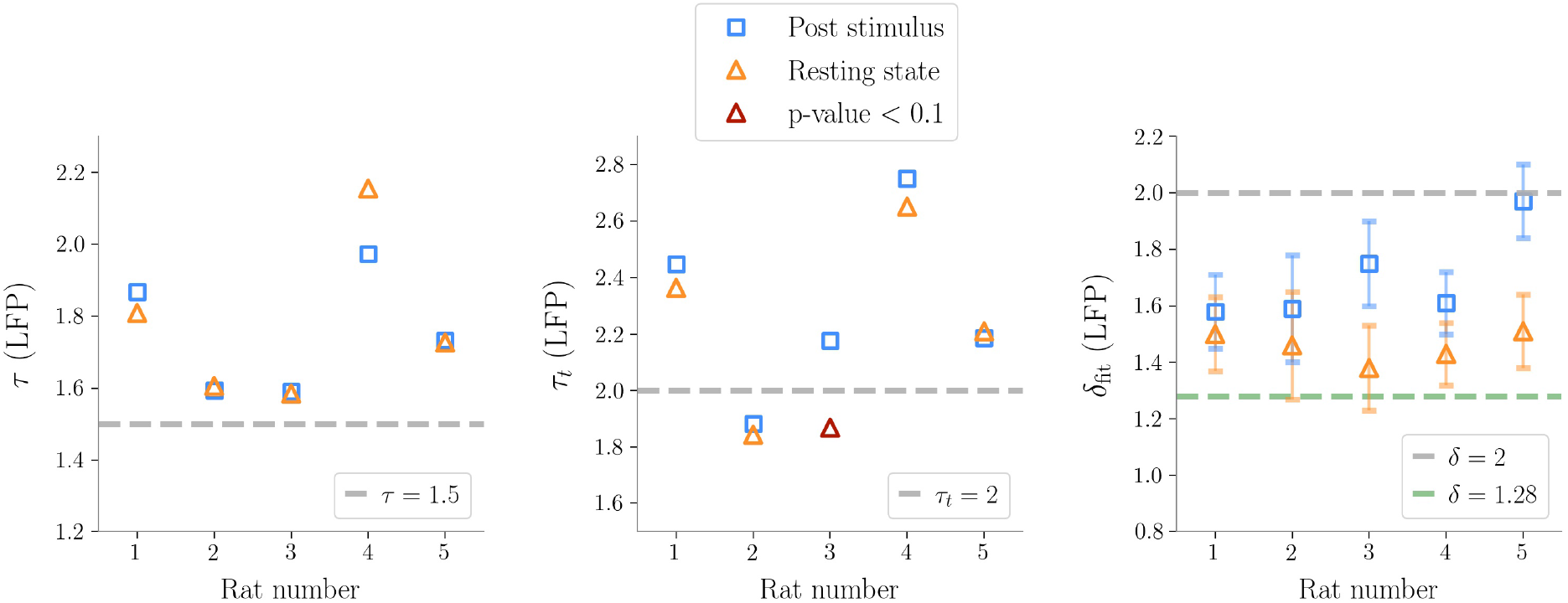
Exponents obtained after subsampling in Local Field Potential data (*τ* is the exponent for the sizes, *τ*_*t*_ for the durations, *δ*_fit_ is the fitted exponent from 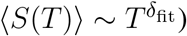. The exponents predicted by the critical branching process are shown in the plot as gray dashed lines. The line *δ* = 1.28 is also plotted, which is reported in [24] as a universal exponent found in many different experiments. It is noteworthy that, as seen in the plot on the right, the exponents *δ*_fit_ are always far from the hallmark of the branching process, except for one case after stimulation, where the presence of large bumps in the size distribution makes the slope of the line in the (⟨*S*(*T*)⟩, *T*) plane steeper.

Then, we analyzed stimulus-evoked responses over two seconds after the stimulus and confirmed power law scaling (accepted by statistical tests). However, at a more careful look, the size distribution was altered by the presence of a *bump* (also known as heap [25]), Figure 3. Clearly, the bump derived from an excess of large size avalanches (i.e., involving a large number of microelectrodes). As suggested by Figure 2, these avalanches were not only large, but also composed by highly synchronous events across cortical layers. Correspondingly, the bump vanished in the duration distribution, confirming that the increase in avalanche sizes was not accompanied by a corresponding increase of lifetimes and the events remained concentrated within a few time bins.

In general, we found that the crackling noise relation was verified both at resting state and post-stimulus (Figure 3 and see Supplementary material for the avalanches results on all the rats). Nevertheless, after the stimulus, a localized deviation from the expected trend was observed in correspondence of the bump found in the size distribution and therefore also attributed to large and synchronized waves of activity unleashed by whisker deflection. As anticipated previously, avalanches durations, and consequently also ⟨*S*⟩ (*T*), extended over about two orders of magnitude. In fact, since LFPs are average signals that integrate over space and time single neuron events, it is expected that the maximum size of the avalanches extracted from LFPs is of the order of the array size (see Supplementary material for an analysis of finite size effects in LFPs avalanches) [6]. Specifically, a cutoff in *P*(*S*) around a value *N*_*C*_ *≈N*_*E*_, with *N*_*E*_ the number of the recording electrodes, is commonly observed experimentally, meaning that during an avalanche each electrode is typically activated just once. This cutoff in *P*(*S*) implies that ⟨*S*⟩ (*T*) *< N*_*C*_, and, from ⟨*S*⟨ (*T*) *∼ T*^*δ*^, it also implies 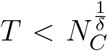 [20]. Thus, if *δ* > 1, the cut-off in *P*(*S*) causes a much earlier cut-off in both *P*(*T*) and ⟨*S*⟩ (*T*).

### 3.3 Avalanches in Spikes Data

Spikes avalanches were analyzed for four rats. Albeit grossly similar to LFPs, results unveiled important differences, first of all the fact that they were almost not affected by the finite size of the recording array. In fact, despite the low number of microelectrodes (twenty-seven sites spanning across the cortex in the vertical direction), avalanches could be observed with size greater than 10^2^ events. The reason is that, in spikes, the same avalanche can reach an electrode repeatedly and in quick succession, contrary to LFPs where single neuron contributions are integrated over space and time within the brain tissue and coalesce to generate the recorded signal [20].

Similarly to LFPs, we found a power law distribution of avalanches both in terms of size and duration, except for one rat deviating from this general trend with respect to duration alone (Table 2 and Figure 6). Moreover, we confirmed the emergence of the bump of activity in the post-stimulus size distribution caused by the abundant number of large-sized avalanches (Figure 5).

**Figure 5.**
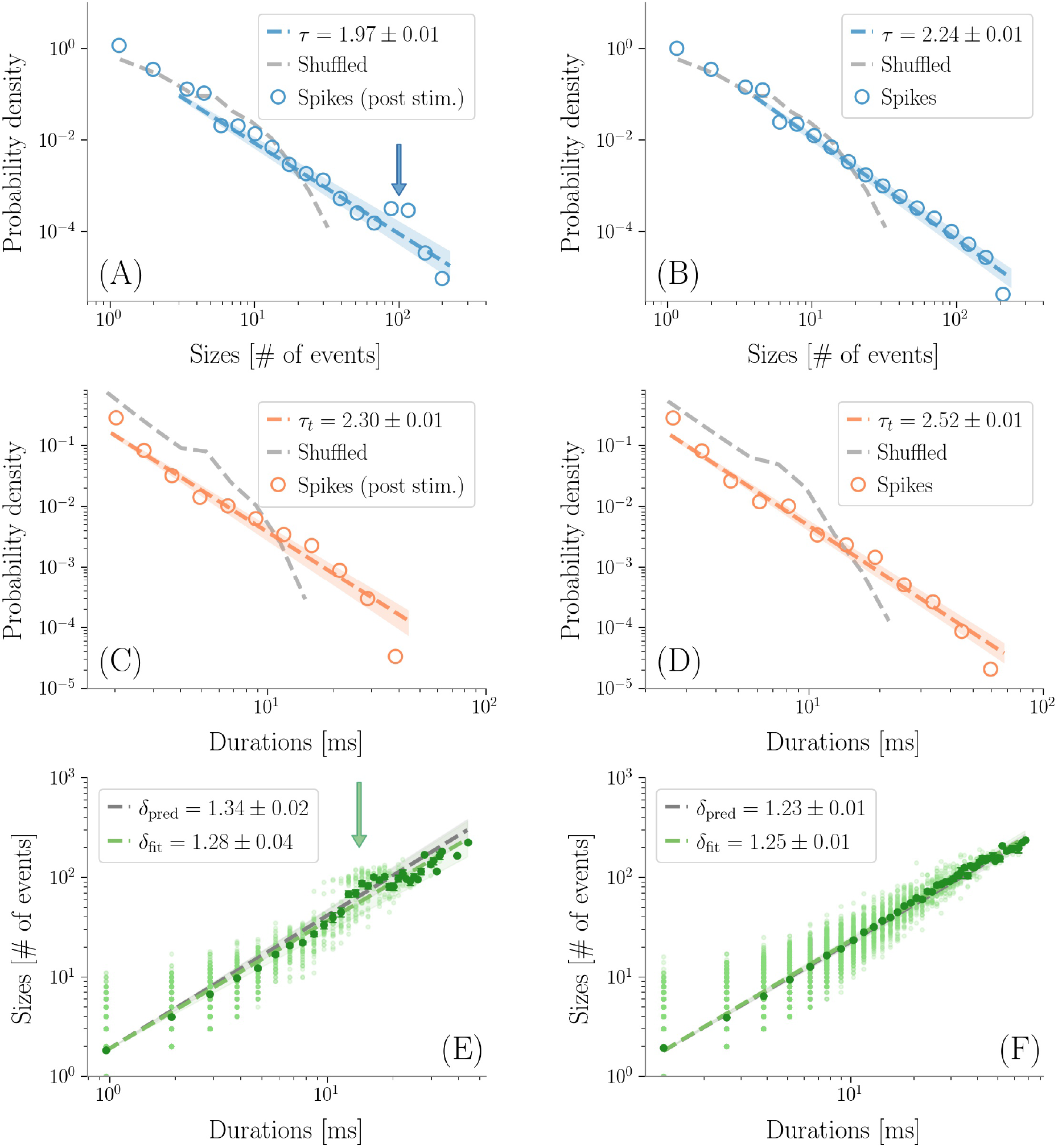
Avalanches in spikes data. Avalanches sizes probability density in one rat after a stimulus (A), and during resting state (B). Notice the bump in the post-stimulus distribution. (C-D) The same, for avalanche durations. Durations are expressed in *ms* by multiplying the number of bins times ⟨IEI⟩. In the randomized datasets the exponential distribution provides a better fit in all cases. The shuffling procedure consists in randomizing the occurrence times of the events of each channel, so that the events rate of each channel is preserved. (E-F) The crackling noise relation is verified in both cases within the experimental errors. Once more, notice in (E) the presence of a bump in the post-stimulus regime.

**Figure 6.**
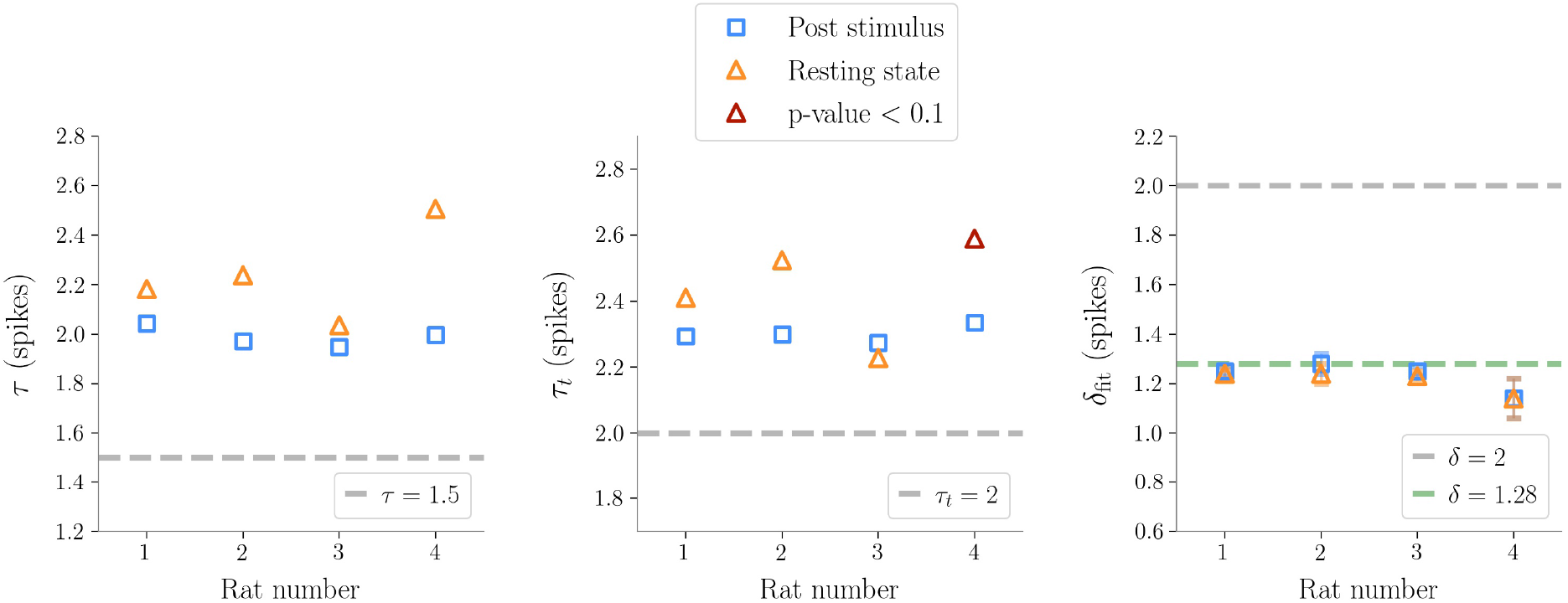
Exponents obtained after subsampling in spikes data (*τ* exponent for the sizes, *τ*_*t*_ for the durations, *δ*_fit_ the fitted exponents of the crackling-noise relation). The exponents predicted by the critical branching process are also plotted. The line *δ*_fit_ = 1.28 is also plotted, which is found in [24], and which turns out to be remarkably close to our results.

The exponents *τ* and *τ*_*t*_ were greater than the ones found in LFPs (Table 2), but also greater than the ones predicted for a critical branching process (Figure 6). Interestingly, the exponent *δ* was consistently close to the value *δ ≈* 1.28 which was found in [24] to be universal, i.e. to hold across different experimental conditions, from cultured slices to freely moving or anesthetized mammals.

Finally, as avalanche size and duration were less affected by finite size effects with respect to LFPs, spikes allowed us to reliably test the crackling noise relation, which was verified in all dataset (Figure 5, see also Supplementary materials). In particular, from the results on spikes data, that are believed to be more robust in this respect [37], we concluded that post-stimulus avalanches statistics is also compatible with the results recently reported by Fontenele and collaborators [24]. Nevertheless, also in this case, the waves of large and synchronized activity triggered by whisker deflection generated a local bump deviating from the crackling noise relation.

## 4 DISCUSSION AND CONCLUSIONS

In this work we investigated criticality in the rat barrel cortex, which offers several advantages over other sensory systems. First, there is a clear and well characterized somatotopic representation of the whiskers in this primary sensory cortex, where single whiskers are basically mapped to single cortical columns with a one-to-one correspondence. This differentiates the barrel cortex from, e.g., the primary visual cortex where the spatial mapping of the inputs does not follow a simple vertical columnar organization.

Second, whisker receptors, which are transducers placed around the follicle, directly activate primary sensory neurons, that therefore encode whisker deflections without interposed processing. In other circumstances, such as in the visual system, the transduced stimulus is subject to extensive processing already at the periphery (e.g., by the retina network) which makes it difficult to disentangle the dynamics and the contributions of cortical and pre-cortical networks in response to sensory inputs. Moreover, the processing pipeline of the whisker somatosensory system is relatively simple and well characterized in mammals, with the trigeminal brainstem nuclei and the thalamus being the only two intermediate stages before the cortex [41, 42]. Third, single-whisker deflection can be controlled with high accuracy through closed-loop piezoelectric systems enabling a tight experimental control over delivered repetitions of sensory stimuli.

As for other sensory cortical areas, it can be hypothesized that the barrel cortex takes advantage of criticality to efficiently map tactile stimuli. This holds also for the single barrel column that faces the severe challenge to represent the parameters related to deflection of its corresponding whisker (e.g., amplitude, direction and velocity of displacement) as transduced by the follicle receptors, in an efficient, noise-tolerant manner and in real-time. According to a simplified general model of the processing in the barrel column, it is believed that layer IV acts as main input stage of sensory information propagating from the thalamus, whereas layer V is the main output. However, the few thousands neurons composing a single barrel form complex microcircuits of excitatory and inhibitory connections across layers. These microcircuits generate a rich dynamics that has been shown, both in-vitro and in-vivo, to include avalanches or synchronization states, such as UP and DOWN states and oscillations, and whose significance in terms of tactile information mapping and processing is far from being understood [43, 44].

We run our investigation in an anesthetized condition (tiletamine) which, contrary to the awake animal, allowed us to precisely control whisker deflection by a piezoelectric actuator. With this anesthetic, spontaneous activity remains rich and contains periods of oscillations and UP and DOWN states, while the activity evoked by whisker stimulation shows clear similarities with the response of the awake animal. In the context of our study, another advantage of the anesthetized animal with respect to awake conditions was that, by reducing cortico-cortical communication, the anesthetic was insulating the somatosensory barrel region from external ‘contaminating’ the waves of activity propagating from other brain areas [29, 45, 46].

We measured neural activity across the six cortical layers of a single barrel both in the domain of spikes and LFPs, thus extending the avalanches analysis over a wide frequency range and covering both single neuron and population dynamics. For LFPs, we used a high-density 2D array of microelectrodes developed at the purpose to monitor at high spatial resolution the electrical potential within a planar section of the barrel column [47]. Contrary to previous work based on imaging methods [48], and in analogy to findings on spikes and LFPs [9, 18], we found signatures of criticality during spontaneous activity also in the anesthetized animal. Avalanche sizes and lifetimes followed power laws during basal activity, both in the case of spikes and of LFPs. Instead, after the stimulus power laws displayed localized bump-like alterations pointing at the emergence of high synchronization across layers.

Although we did observe inter-rats variability in the power-law exponents (Fig. 4 and 6), the exponent of the crackling noise relation consistently converged to *δ ≈* 1.28, hinting at a critical behavior of the neural network activity across the cortical barrel column at rest [24] (Fig. 6). On the other hand, the spectrum of alternative hypotheses is broad. As recently pointed out [49, 20], this value may also emerge as a spurious result when subsampling spikes data in presence of an underlying process belonging to the branching process universality class. Moreover, different models with different phase transitions [15, 50, 25] may yield non-trivial exponents compatible with the ones found here. In general, the exponents we obtained are anomalous – in the sense that they differ for what reported in awake mammals such as, e.g., in monkey [7] – both for spikes and LFPs. A possible reason is the action of the anesthetic drug tiletamine, as similar deviations were reported previously [9].

At the same time, it is of the utmost importance to note that, with respect to the generative mechanism of the observed avalanches and exponents, it is not clear how a critical branching process could describe our experimental results. First, it assumes a clear separation of time scales between avalanches, and refers to slowly driven systems [51]; the barrel cortex, instead, receives continuous inputs from the thalamus, in addition to other cortical areas such as the secondary somatosensory cortex and the motor cortex. Second, it has still to be clarified how *mean field* exponents can arise in biologically realistic, non trivial networks [52, 53], and in the presence of feed-back loops such as the ones intrinsic to the barrel cortex. A number of feed-back connections within and among layers are present, and the connectivity is therefore remarkably different from the feed-forward one that was originally assumed to support the critical branching process [6]. We also note that generative mechanisms with absorbing states, such as the branching process, implies no correlation between avalanches [15]. However, in our data, we found non-negligible values of *τ* ^*\**^ hinting at the existence of temporal correlations between subsequent avalanches. Our results, on the other hand, are thus in agreement with previous works in which long range temporal correlations emerged through a detrended fluctuations analysis (DFA) of the signals [18, 54, 55]. Although the nature of such correlations remains unknown, we speculate that they may derive from a variety of neuronal and circuital mechanisms including, e.g., a neuromodulation of the barrel column, acting at time scales that are large compared to avalanches lifetimes.

Moreover, in the light of carefully investigating the underlying phase transition, avalanches characterization is sensitive to several parameters: exponents depend on the temporal bin chosen, and sampling effects may definitely bias the avalanches distribution. The first aspect is linked to the fact that, in absence of a clear separation of time-scales, it is not guaranteed that the avalanches found with the method of temporal binning reveal the underlying causal avalanches. For example, avalanches revealed by a time binning do not reproduce causal avalanches if they initiate simultaneously, or in the case of high stimulation rate [19, 56]. Recent works are indeed showing that both LFPs and spikes based avalanches may be problematic: in the former case, they may not be able to distinguish between a critical, a subcritical or supercritical phase [20, 23]; in the latter, it has been shown that the exponents of the distributions appear to be distorted [20, 49]. Recently, some first attempts were made to characterize criticality in neural systems in a broader sense [57, 58, 59], but as of now, and despite all these unsolved issues, power law avalanches are still the most employed approach to test the critical brain hypothesis.

Nonetheless, let us note that in our work we consistently find bumps in the avalanche sizes distributions in the periods after stimuli. These bumps mark avalanches with large sizes that appear with a higher probability than what would be predicted by the power law scaling - in the case of LFPs, these sizes are of the order of the array size. This interesting but seemingly simple observation has quite profound implications. In fact, the bumps are clearly related to large synchronization events that take place after stimuli, with avalanches characterized by highly synchronous activity across microelectrodes (see Fig. 2). These avalanches are related to the strong response to the stimulus that emerges prominently first in layer IV and then quickly spreads among the other layers, along a prevalent vertical direction, thus eliciting a global activity in the barrel. Therefore, the presence of the bumps is clearly related to the synchronization behavior of the barrel cortex and thus suggests that synchronization waves and oscillations play a fundamental role in shaping the neural activity during a time window dedicated to map the tactile stimulus in the brain. In recent years, some efforts in this direction were made [24, 25] but a comprehensive model of this kind of transition remains elusive.

In conclusion, on the one hand we believe that, in future works, it will be important to include other markers of criticality in the analysis that go beyond neuronal avalanches. For instance, we used the same LFP data to study the spatial correlations in the resting state and look for long-range correlations that are an hallmark of criticality [60], but one could also consider patterns of dynamic functional connectivity, e.g. obtained by computing the instantaneous phase difference between signals at different locations [61]. These kind of analysis should be extended to include the behavior during and after the sensory input, going beyond the usual resting state studies. On the other hand we also believe that sound models of the possible self-organizing mechanisms of the cortex will have to necessarily take into account its oscillatory behavior, whose signature in the present work is the emergence of the bump in the avalanche size distribution due to the major synchronizations within the cortical network. In perspective, avalanches and oscillations may represent two faces of the cortical brain dynamics medal, two intertwined and functionally relevant processes that will have to be dealt with together in future theoretical and experimental investigations on the sensory coding in the brain.

## AUTHOR CONTRIBUTIONS

SV and SS designed the study. SV, SS and MB supervised the research. MM conducted the experiments. RO designed the whisker control system. BM analyzed the data. BM and GN prepared the figures. BM, GN, MB, MM, SS and SV interpreted the results and wrote the article. All authors contributed to the article and approved the submitted version.

## ACKNOWLEDGMENTS

We thank A. Maritan for fruitful discussion.

## DATA AVAILABILITY STATEMENT

The dataset generated for this study is available upon request to

## SUPPLEMENTARY MATERIAL

**Figure S1.**
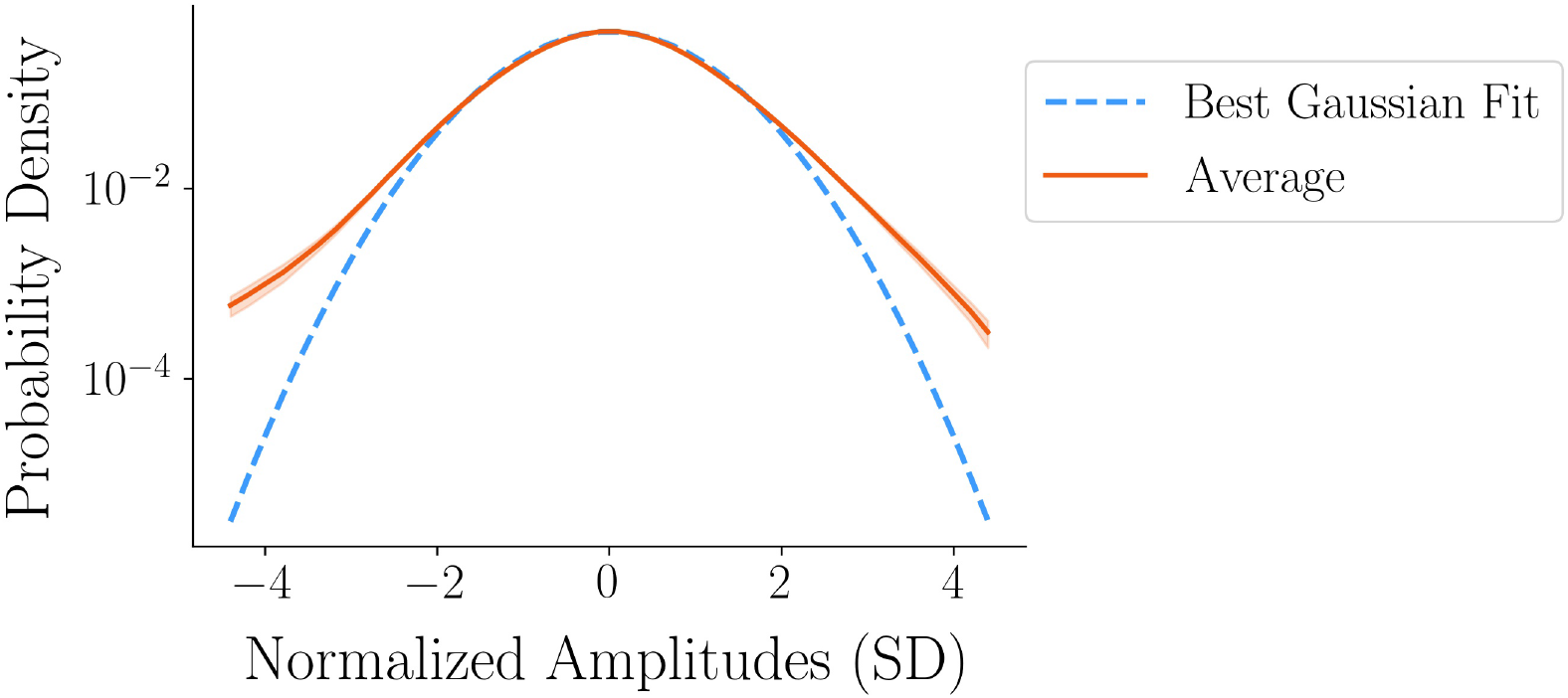
LFPs events threshold definition. The red curve depicts the grand average of the signals amplitude distributions over all channels and trials in LFPs of one rat. Note that the signal from each channel is z-normalized by subtracting its mean and dividing by the SD. The dashed line depicts the best fit of a Gaussian distribution to the data for the range between + 4.5 SD and - 4.5 SD. The Gaussian fit starts deviating from the average signal at around ± 2 SD. Hence, in order to avoid false positives, we set the event threshold at ± 3 SD. A logarithmic scale is used for the *y*-axis.

**Figure S2.**
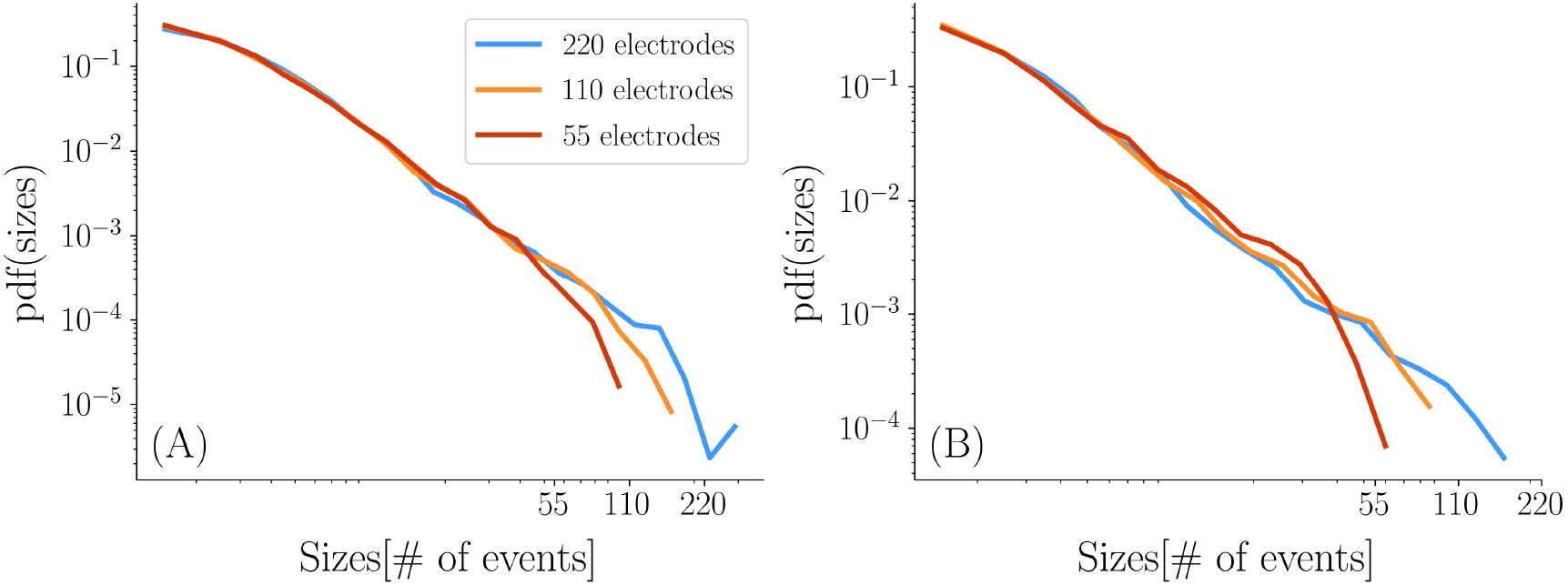
Power law finite size effects analysis for LFP avalanche sizes at resting state. We verify that the cutoff in LFP avalanches distribution is dependent on the size of the array. For this purpose, we repeat the avalanche analysis considering only halves and quarters of the array. For example, when considering quarters, only single columns of the array 55 *×* 4 are considered in the analysis: the array is split along the direction of the barrel column, in order not to create halves/quartets with different behaviors due to the inclusion of different layers. The results from the four columns are averaged to produce the analysis for a quarter of the array. The same procedure is applied to the two halves of the array. We verify that the maximum size of the avalanches (called here *N*_*C*_) is dependent on the number of electrodes (*N*_*E*_) of the array. It results that *N*_*C*_*≥ N*_*E*_ when considering both positive and negative excursions of the signals as events (Fig. S2 A), as in the main text [26], and it results that *N*_*C*_ ≈ *N*_*E*_ when only negative excursions are considered as events (Fig. S2 B). As noted in the main text, this cutoff in the avalanche sizes induces a much earlier cutoff in avalanche durations. Also, the exponents do not change when reducing the number of electrodes.

**Figure S3.**
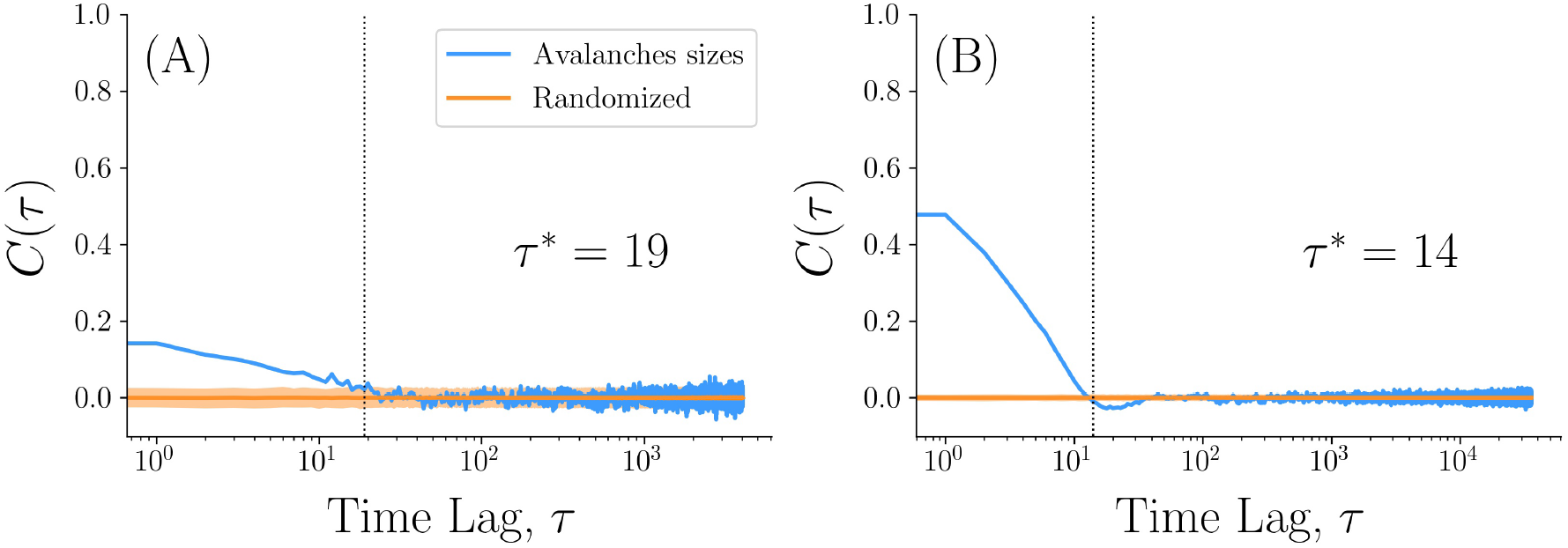
Are avalanches correlated? As these plots show, subsequent avalanches are correlated. The autocorrelation function of the logarithm of avalanche sizes during resting state is computed following [31]. (A) Avalanches in LFPs data display a characteristic autocorrelation time of *τ*^*\**^ = 19, i.e. on average *τ*^*\**^ consecutive avalanches are correlated. (B) In spikes data, *τ*^*\**^ = 14.

**Figure S4.**
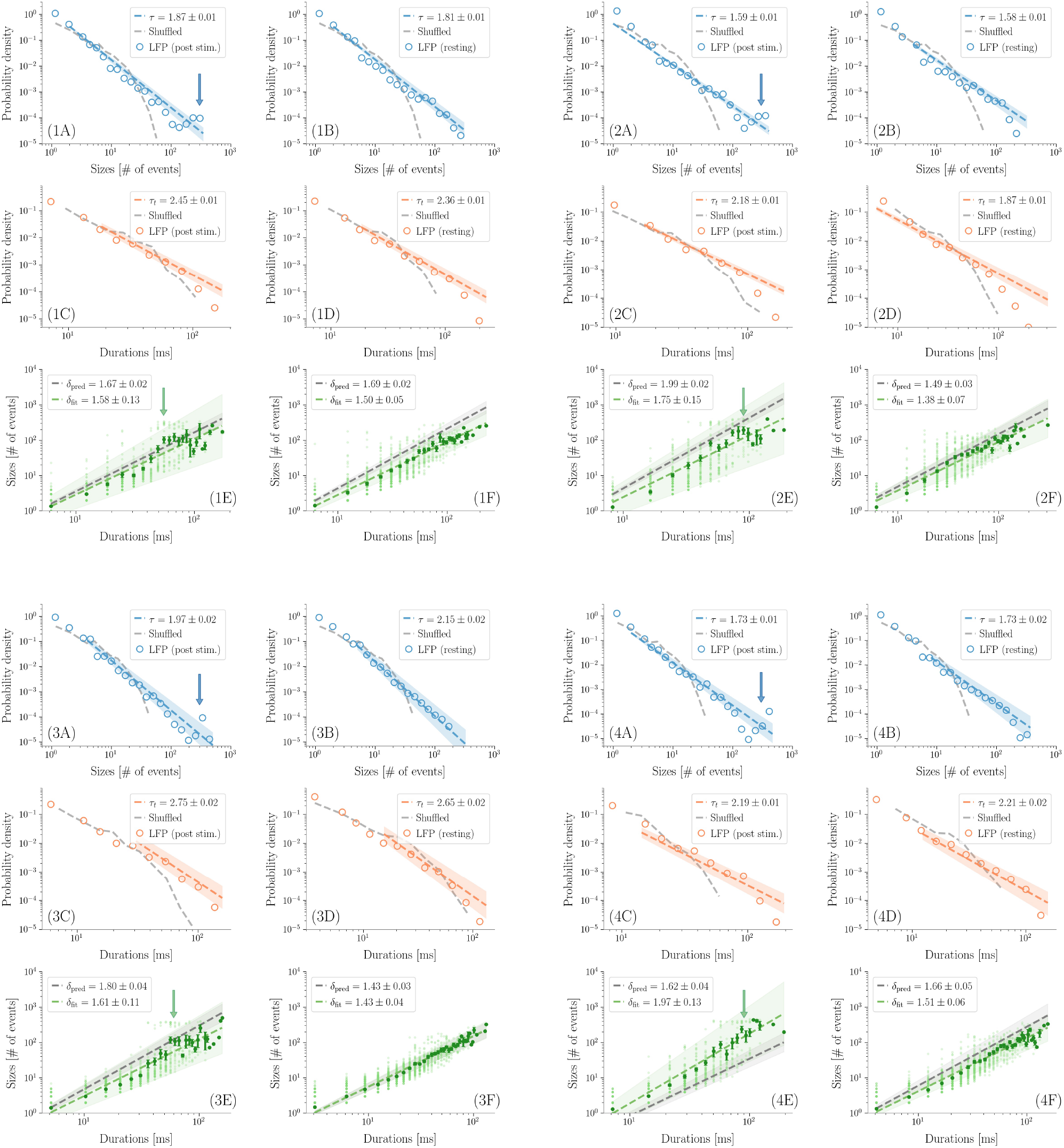
Distribution of avalanche sizes (in blue), durations (in red) and crackling noise relation (in green) in LFPs data obtained as described in the main text, both at resting and post stimulus for four different rates. Detailed avalanche statistics results are presented for all rats in the main text.

**Figure S5.**
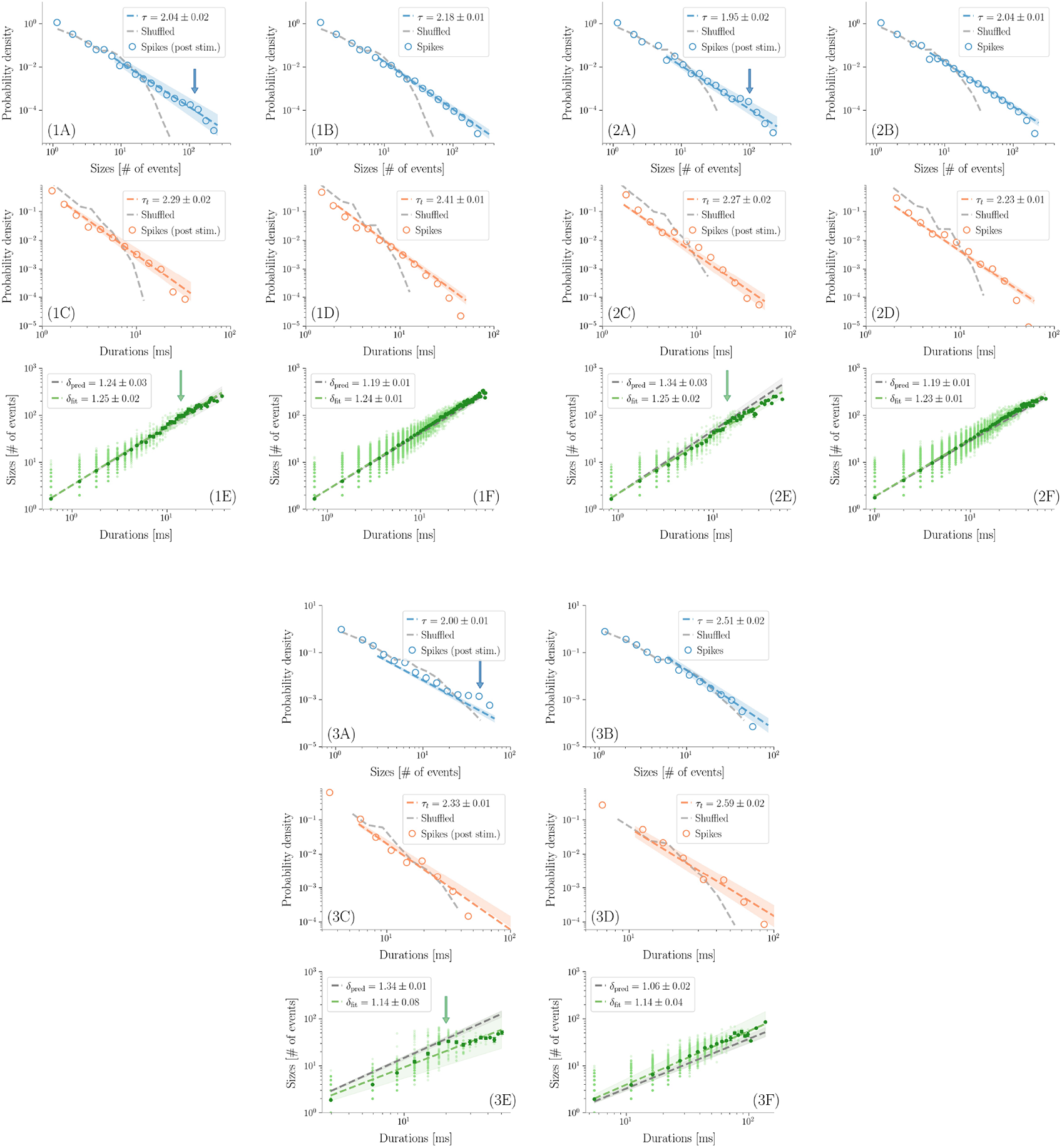
Distribution of avalanche sizes (in blue), durations (in red) and crackling noise relation (in green) in spikes data obtained as described in the main text, both at resting and post stimulus for three different rats. Detailed avalanche statistics results are presented for all rats in the main text.

